# sTAM: An Online Tool for the Discovery of miRNA-set Level Disease Biomarkers

**DOI:** 10.1101/2020.04.10.035345

**Authors:** Jiangcheng Shi, Qinghua Cui

## Abstract

microRNAs (miRNAs) are one class of important small noncoding RNA molecules, which have shown their excellent ability as biomarkers of various diseases. However, current miRNA biomarkers including those comprised of multiple miRNAs work at a single-miRNA level but not at a miRNA set level. Given the rapidly accumulated miRNA omics data, it is believed that miRNA set level analysis could be an important supplement to the single miRNA level analysis. For doing so, here we presented a computational method for single-sample miRNA set enrichment analysis and developed the sTAM tool (http://mir.rnanut.net/stam). Moreover, we demonstrated the usefulness of sTAM scores in discovering miRNA-set level biomarkers through two case studies. We conducted pan-cancer analysis of the sTAM scores of “tumor suppressor miRNA set” on 15 types of cancers from TCGA and 14 types of cancers from GEO, finding that the scores show a good performance in distinguishing the cancers from the controls. Moreover, we revealed that the sTAM score of “brain development” miRNA set can effectively predict cerebrovascular disorder (CVD). Finally, we believe sTAM can be used in discovering disease-related biomarkers at a miRNA-set level.

## Introduction

MiRNAs are one class of important small non-coding RNA molecules with a remarkable variety of biological functions.^1^ Recently, miRNAs have been proposed as being useful in diagnostics and prognosis as biomarkers for a series of diseases.^2^ However, as a matter of fact, most of the current miRNA-based biomarkers work at a single miRNA level. As is known to all, no molecules work in isolated but have various connections with others, and enrichment analysis represents one important and popular method to infer the relationships. In addition, single sample gene set enrichment analysis (ssGSEA) stands for a powerful tool to find gene set level biomarker for various diseases.^3, 4^ And gene set level biomarker represents an important supplement to single gene level biomarker. However, methods and tools for single sample miRNA set enrichment analysis (MSEA) are still not available.

For doing so, we presented sTAM and implemented the web server. Moreover, by applying sTAM to 29 cancer miRNA expression datasets from TCGA and GEO, we showed that the sTAM score of tumor suppressor miRNA set^5^ can discriminate tumors from controls (healthy or adjacent tissues) well. In addition, using sTAM, brain development miRNA set was identified as an effective biomarker for predicting CVD. These result suggest that sTAM scores of various miRNA sets have the potential to be biomarkers for monitoring disease formation and development.

## Results

### Overview of sTAM server

The interface of sTAM was shown in Figure 1. sTAM works according to the following procedure. At first, users are required to upload the whole genome-wide miRNA expression profile. Then, users need to select a reference miRNA set or input their own annotated miRNA set instead. In addition, users can adjust advanced parameters, for instance, filter out too small or too large miRNA sets. Finally, after running sTAM, users will get the sTAM score of each inputted sample on each miRNA set. And with these scores, users can conduct further specific analyses.

**Figure 1.**
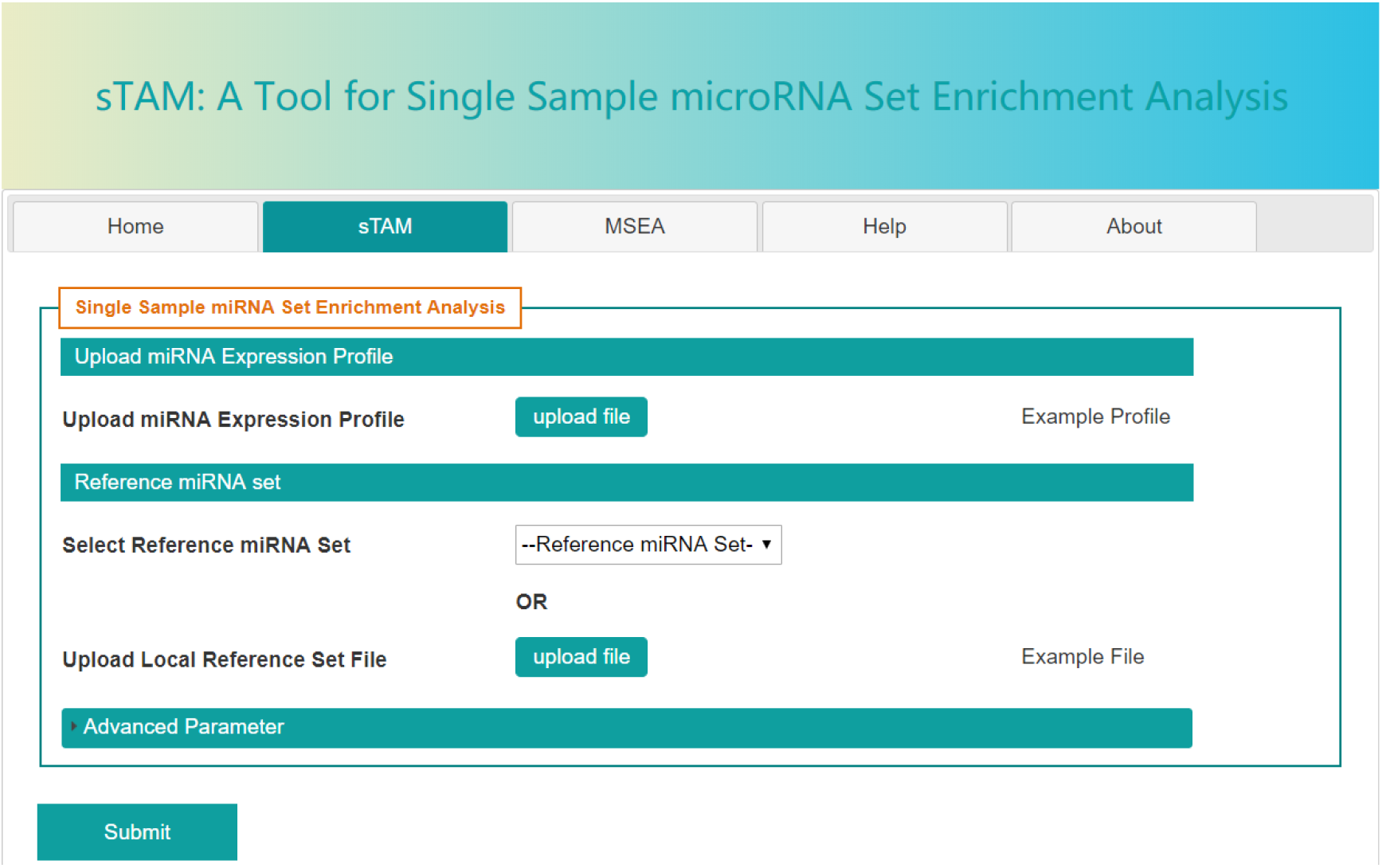
The interface of sTAM.

### Case studies

#### Case 1: sTAM score in tumor suppressor miRNA set effectively discriminates cancers from controls

It is well known that some miRNAs are tumor suppressors, which play critical roles in cancer formation and development. It is thus hypothesized that the sTAM score of the tumor suppressor miRNA set in miRNA expression profiles could discriminate cancers from normal controls. To confirm this hypothesis, we then applied the sTAM tool to 15 datasets from TCGA (https://cancergenome.nih.gov/) and 14 datasets from GEO (datasets with at least 50 samples, https://www.ncbi.nlm.nih.gov/geo/). As a result, the sTAM score of tumor suppressor miRNA set has a good performance for discriminating cancer from normal controls for most of the datasets from TCGA (Figure 2A) and for most of the datasets from GEO (Figure 2B).

**Figure 2.**
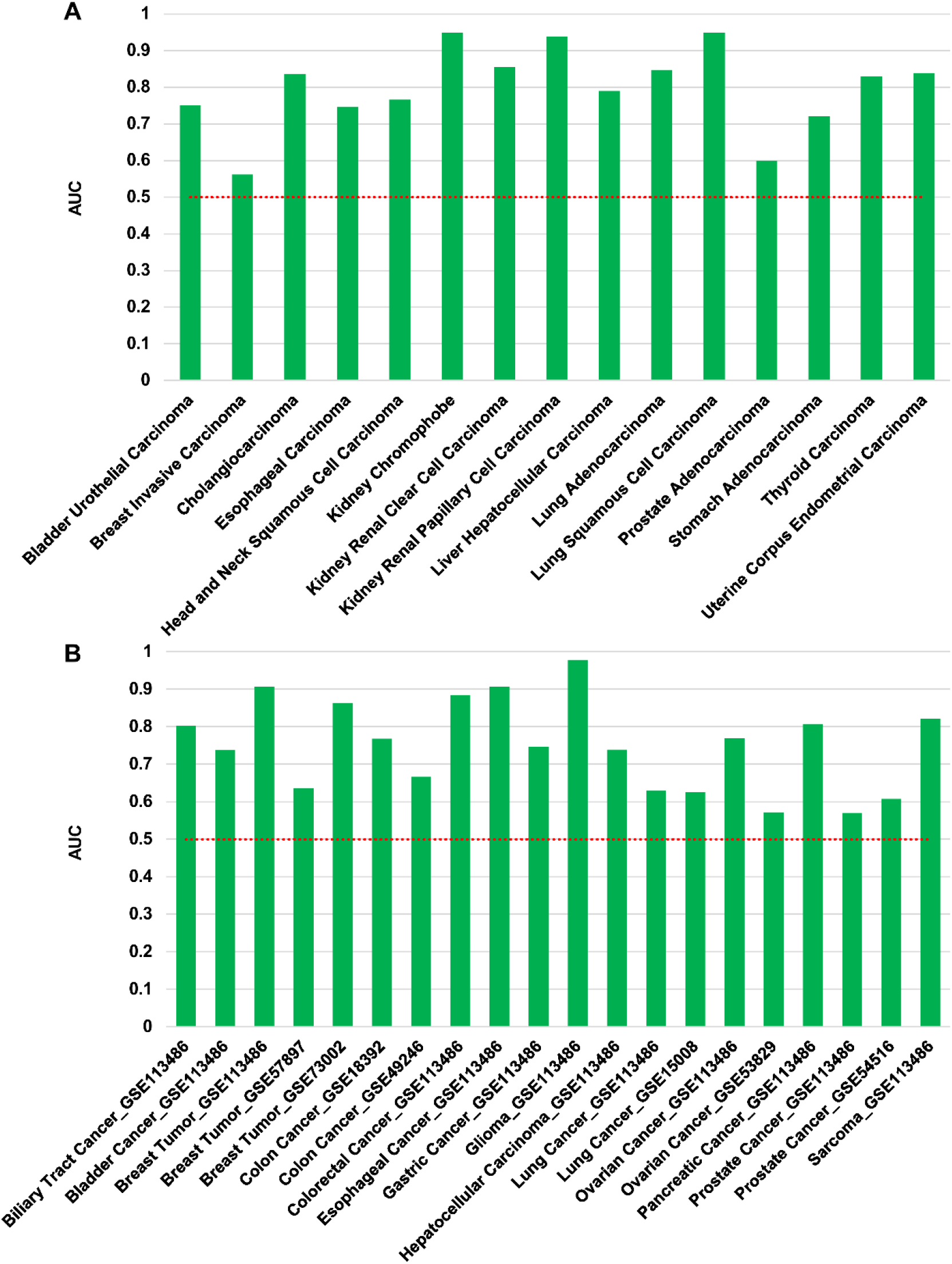
AUCs for the sTAM score of tumor suppressor miRNA set in discriminating cancers from normal controls in datasets from TCGA (A) and datasets from GEO (B). The height of each bar represents the AUC in corresponding dataset.

#### Case 2: sTAM score in brain development miRNA set effectively predicts CVD

There exist a large number of miRNAs in body fluids such as blood, urine and saliva.^6^ Thus, circulating miRNAs can be used as potentially valuable disease-related biomarkers. CVD is one most common and severe complex disease but effective biomarkers are still limited. Here we ask whether sTAM strategy can predict CVD. Given that CVD is highly associated the process of brain development, we investigated the sTAM scores’ ability for predicting CVD, in order to verify whether the sTAM scores of the brain development set are different between patients with CVD and non-CVD controls, and further confirm the biomarkers of miRNA-set level related to CVD. For doing so, we obtained the serum miRNA expression profiles of CVD patients and non-CVD controls from GEO, and performed sTAM analysis on the brain development miRNA set. As a result, we found that there are significant differences when comparing the sTAM scores of brain development miRNA set of CVD patients with non-CVD controls (Figure 3A, *P*-value = 2.88e-44, Wilcoxon rank-sum test). In addition, the sTAM score of brain development miRNA set achieves an AUC of 0.822 for discriminating CVD patients from non-CVD controls (Figure 3B), suggesting that the miRNAs in the brain development set could have potential as biomarkers for CVD prediction at miRNA set level.

**Figure 3.**
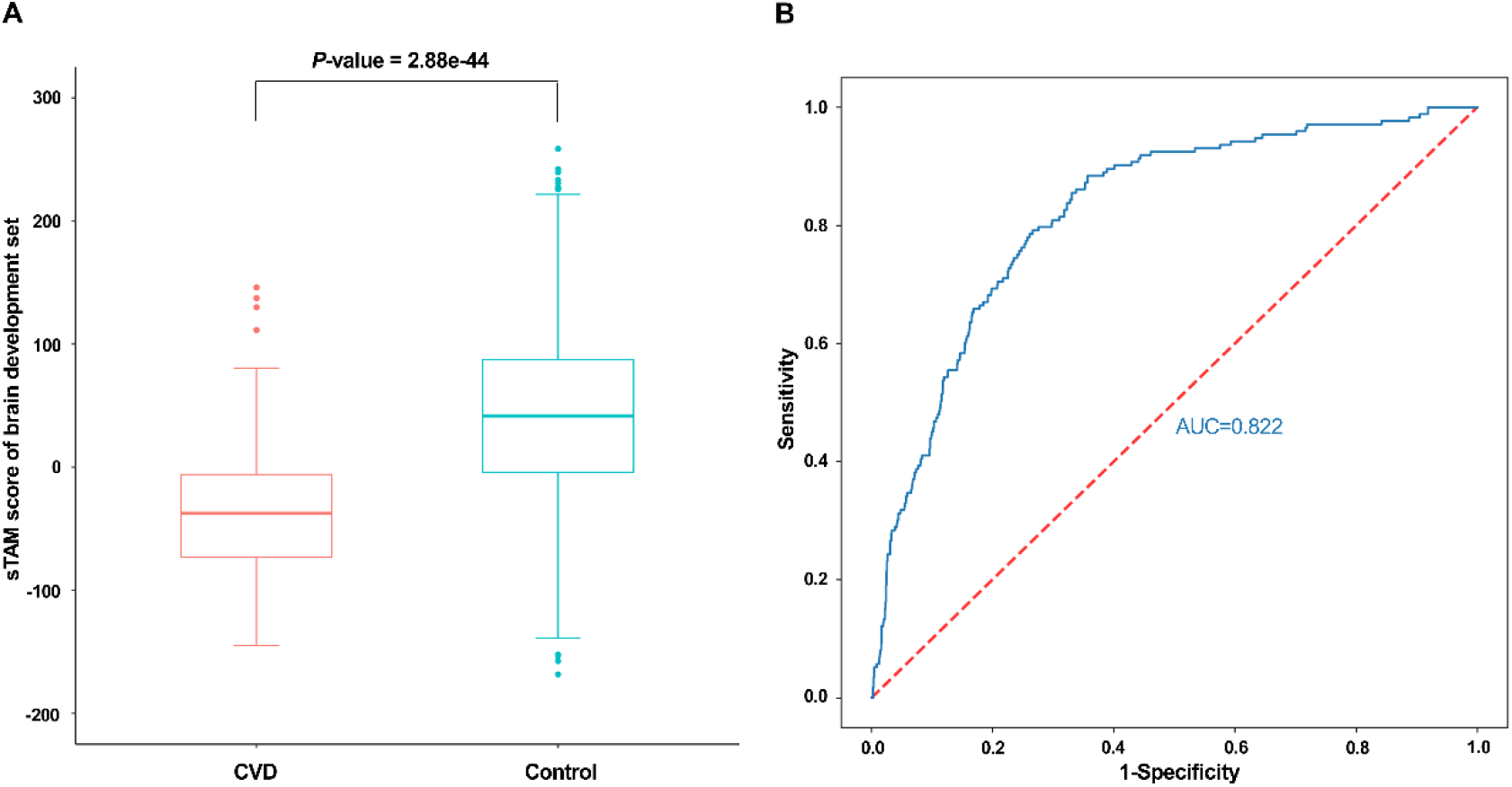
Comparison of sTAM score of brain development miRNA set between CVD patients and healthy controls (A) and the AUC in discriminating CVD patients from non-CVD controls (B).

## Discussion

MiRNAs are one class of important small non-coding RNAs that mainly involved in the post-transcriptional regulation of gene expression and play crucial roles in a series of biological processes and various disease development, for example cancer. Recently, miRNAs have shown their excellent ability as biomarkers for various cancer and other diseases.^7, 8^ Given that lots of miRNAs were identified in a number of body fluids and miRNAs normally show high stability,^9, 10^ which makes them suitable to be biomarkers for many diseases. However, the current miRNA-based biomarkers work at a single miRNA level. Even for biomarkers composed of multiple miRNAs, each miRNA represents one component of the biomarker but not work at the level of miRNA set.

We previously defined the concept of miRNA set and found that disease often associated with miRNAs at miRNA set level.^11^ We then developed TAM for MSEA for a list of miRNAs, for example deregulated miRNA in some cancers.^12, 13^ In addition, two other tools for MSEA, miSEA^14^ and miEAA,^15^ have also been developed with different statistical methods. These tools provide helps in discovering valuable clues and predicting novel disease-related miRNAs based on miRNA expression data. However, none of them can perform enrichment analysis at single-sample level, which is very important miRNA-set based biomarker discovery.^4^

Here we present a computational method for single sample miRNA set enrichment analysis and developed the software, sTAM. We integrated 1,236 miRNA sets into sTAM. This makes it easy for users to select the reference miRNA sets provided by sTAM. In addition, users could input their own defined miRNA sets as well. Finally, we proposed two case studies that revealed the sTAM score of tumor suppressor miRNA set can effectively discriminate cancers from controls while sTAM score of brain development miRNA set have good performance in distinguishing cerebrovascular disorders from controls. According to our knowledge, sTAM represents the first tool for single sample miRNA set enrichment analysis. With the continuous improvement of miRNA set annotations and rapid accumulation of miRNA omics data, we believe that sTAM would be a useful tool for the discovery of miRNA-set based biomarkers in various diseases.

## Materials & Methods

### Data sources and data preprocessing

The reference miRNA sets were downloaded from TAM 2.0,^12^ which are composed of six categories: function, disease, family, cluster, tissue specific and transcription factor, a total of 1,236 miRNA sets. As for the case studies, We downloaded 15 cancer miRNA expression datasets from The Cancer Genome Atlas (TCGA) database (https://cancergenome.nih.gov/) and 14 cancer miRNA expression datasets from GEO (datasets with at least 50 samples, https://www.ncbi.nlm.nih.gov/geo/), respectively. The “DESeq2” R package was used to estimate expression value and “sva” package to remove batch effects on the TCGA datasets. In addition, we obtained the serum miRNA expression profiles constituted by 173 CVD patients and 1,612 non-CVD controls from the GEO dataset (accession number: GSE117064).

### Workflow of sTAM

The workflow of sTAM was presented in Figure 4. sTAM allowed submission of whole genome-wide miRNA expression profile and no need to do too much data processing by users. Since the reference miRNA sets are at pre-miRNA level, sTAM would convert the mature miRNA names to precursor names and take the average value of the duplicated miRNAs. Then, sTAM will perform single sample miRNA set enrichment analysis in the background. Finally, the server would return the sTAM scores of each sample on each miRNA set to users. In addition, user could download the compressed result file generated by the server and conduct further analyses.

**Figure 4.**
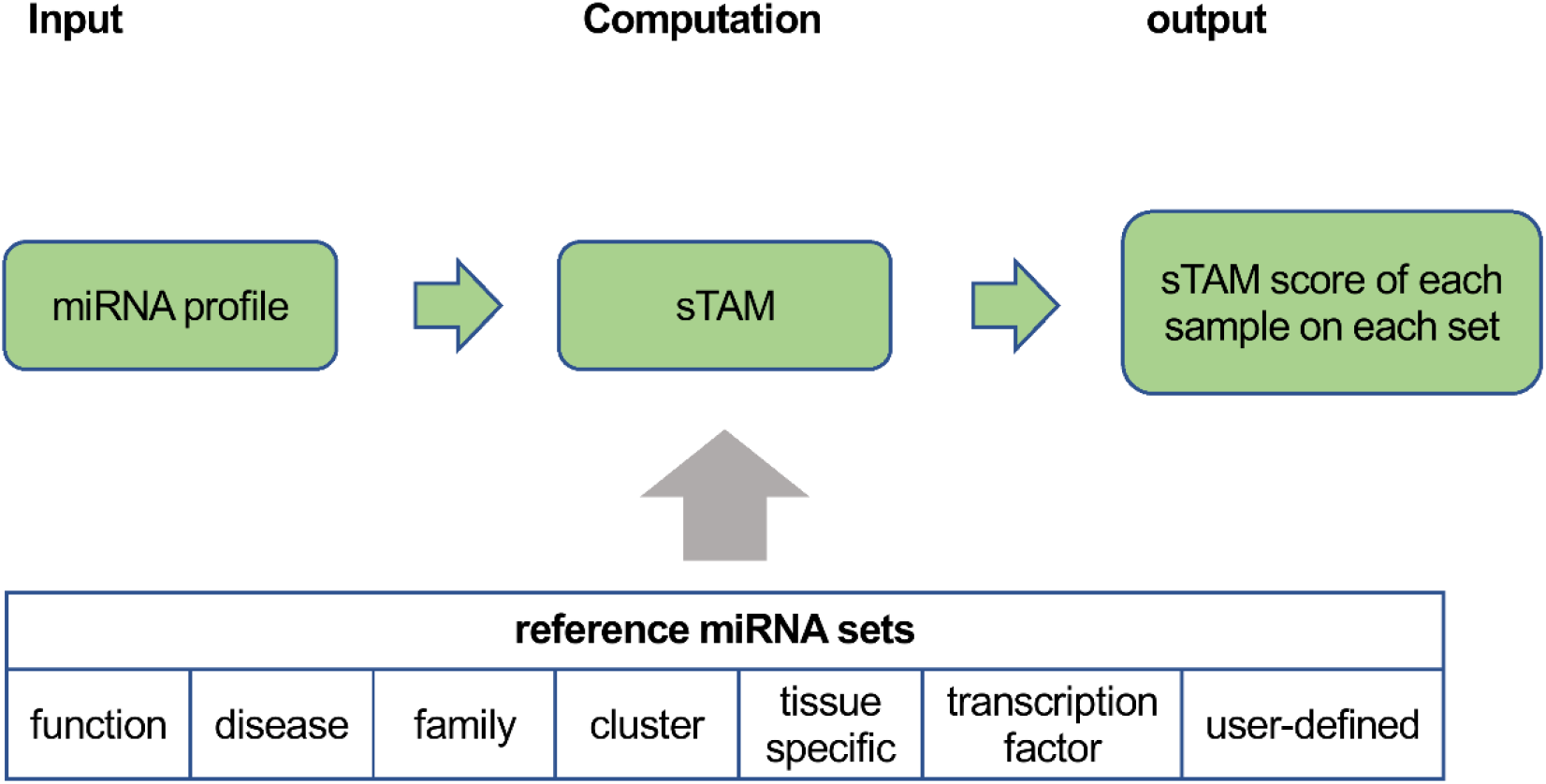
The workflow of sTAM.

### sTAM algorithm

For performing single sample miRNA set enrichment analysis, we adopted the algorithm for single sample gene set enrichment analysis (ssGSEA).^4^ Given a miRNA expression profile, sTAM ranks miRNAs by absolute expression values in one sample and then calculates the enrichment score (ES) by walking down the ranked list of miRNAs, increasing a running-sum statistic when a miRNA is in the miRNA set and decreasing it when it is not. The ES is the maximum deviation from zero encountered in walking the list. In other words, for a given miRNA set *M* of size *N*_*M*_ and single sample *S*, of the dataset of *N* miRNAs, the miRNAs are sorted according to their absolute expression values from high to low: *E* = {*e*_1_, *e*_2_, …, *e*_*N*_}. An enrichment score *ES*(*M*, *S*) is obtained by a sum of the difference between a weighted Empirical Cumulative Distribution Functions (ECDF) of the miRNAs in the miRNA set 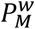 and the ECDF of the remaining miRNAs 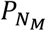:

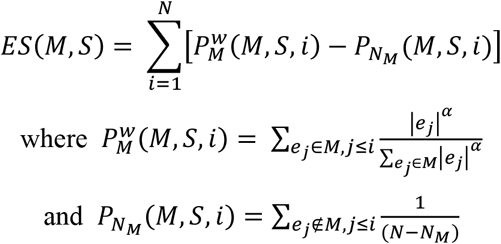

sTAM repeated this calculation for each miRNA set and each sample in the expression dataset. Note that the exponent *α* is set to 0.25 by default based on experience, and adds a modest weight to the rank.

### Server construction

The web server was established in the “Linux + Apache + Thinkphp (version 3.2) framework. The single sample miRNA set enrichment analysis was implemented by using GSEAPY (version 0.9.15), a python wrapper for enrichment analysis. In addition, users are provided with example files and tutorial to get started easily. The sTAM web server is freely accessible at http://mir.rnanut.net/stam

### Statistical analysis

Receiver operating characteristic (ROC) curve and the area under the curve (AUC) were used to express the performance of sTAM score. The Wilcoxon rank-sum test was applied to detect differences of sTAM scores between CVD patients and non-CVD controls by Scipy (version 1.1.0), an open-source scientific computing library for the Python programming language. *P*-value of less than 0.05 was considered statistically significant.

## Author Contributions

Q.H.C conceived the study. J.C.S developed the web server and performed bioinformatics analysis. All authors wrote the manuscript.

## Compliance and ethics

The authors declare that they have no conflict of interest.

## Acknowledgements

This work has been supported by the grants the Natural Science Foundation of China (81970440, 81670462, and 31930056), grants from the Peking University Basic Research Program (BMU2020JC001), Peking University Clinical Scientist Program (BMU2019LCKXJ001), and the Fundamental Research Funds for the Central Universities.

